# Joint single-cell measurements of surface proteins, intracellular proteins and gene expression with icCITE-seq

**DOI:** 10.1101/2025.01.11.632564

**Authors:** Kelvin Y. Chen, Yusuke Takeshima, Andre L. Zorzetto-Fernandes, Katsuhiro Makino, Tatsuya Kibayashi, Mayu Hata, Kenji Ichiyama, Josh Croteau, Kevin Taylor, Shimon Sakaguchi

## Abstract

**The development of single-cell RNA-sequencing assays has transformed our understanding of cellular and tissue heterogeneity, yielding significant insights into disease biology and its underlying mechanisms. In this work, we describe icCITE-seq (intracellular cellular indexing of transcriptomes and epitopes), a scalable method that simultaneously measures surface and intracellular protein levels alongside gene expression across thousands of cells. We validate the specificity of intracellular staining and demonstrate the utility of this multi-omic approach in interrogating phenotypic cellular states through targeted genetic perturbations in primary human T cells. icCITE-seq enables systematic profiling of gene expression, coupled with cytoplasmic, nuclear and PTM epitopes, providing an integrated approach towards understanding cellular identity, complexity and disease regulatory mechanisms.**

## Main

The recent advent of high-throughput technologies enabling comprehensive single-cell measurements has revolutionized our ability to dissect complex cell types and understand cellular states that underlie cellular heterogeneity. While profiling of single modalities has been invaluable for cellular characterization and phenotyping, new techniques that couple multiple modalities into a single assay present new opportunities to gain further insights into the interconnected layers of cellular regulation and function. For example, the concomitant detection of transcriptome and surface proteins in CITE-seq^1,2^ and REAP-seq^3^, which complements sparse mRNA measurements with robust protein quantification, and its extension to genetic perturbations^2,4^ has enabled nuanced cell type annotation and functional assessment. Similarly, paired measurements of chromatin accessibility and protein profiling in ASAP-seq allow for sensitive multiplexed surface and intracellular protein quantification^5^.

While single-cell assays profiling the transcriptome and concomitant surface proteins are well established, accurately quantifying intracellular epitopes remains challenging. Indeed, recent advances in multi-modal single cell assays targeting mRNA and intracellular epitopes are often met with issues over loss of RNA integrity^6,7^ and removal of surface and cytoplasmic epitope fractions^6,8^.

Here, we establish icCITE-seq, a massively parallel protocol for sensitive quantitation of surface and intracellular epitopes, that retains high-quality transcriptome data, and combine it with ASAP-seq to phenotype CRISPR perturbations in human primary T cells. We demonstrate the wide applicability of icCITE-seq in profiling cytoplasmic, nuclear and phosphorylated epitopes, enabling comprehensive characterization of cellular states and the identification of key regulatory mechanisms involved in T cell activation and differentiation.

## Results

### icCITE-Seq couples detection of surface and intracellular epitopes with transcriptomic profiling

To integrate intracellular epitope quantification with single cell transcriptomic profiling, we developed intra<u>c</u>ellular <u>C</u>ellular Indexing of <u>T</u>ranscriptomes and <u>E</u>pitopes (icCITE-seq), which enables the simultaneous quantitation of gene expression with surface and intracellular epitopes. In this method, surface protein immunophenotyping is performed by first staining cells with oligonucleotide-conjugated antibodies (TotalSeqA^TM^), followed by fixation and permeabilization by a methanol-based fixative (**Fig. 1a**). The permeabilized cells are then stained with additional oligonucleotide-conjugated antibodies (TotalSeqB^TM^) targeting intracellular epitopes and then profiled with microfluidic-based scRNA approaches (**Supplementary Note**). To benchmark icCITE-seq and further dissect the molecular machinery governing T cell activation, we applied a combinatorial hashing approach^5^ to profile CRISPR perturbations targeting 107 nuclear factors in primary human CD4 T cells undergoing TCR stimulation with a panel of 277 antibodies targeting surface markers and 40 antibodies targeting intracellular epitopes using ASAP-seq (85,641 cells), CITE-seq (95,805 cells) and icCITE-seq (59,777 cells) (**Extended Data Fig. 1a-c**).

**Figure 1.**
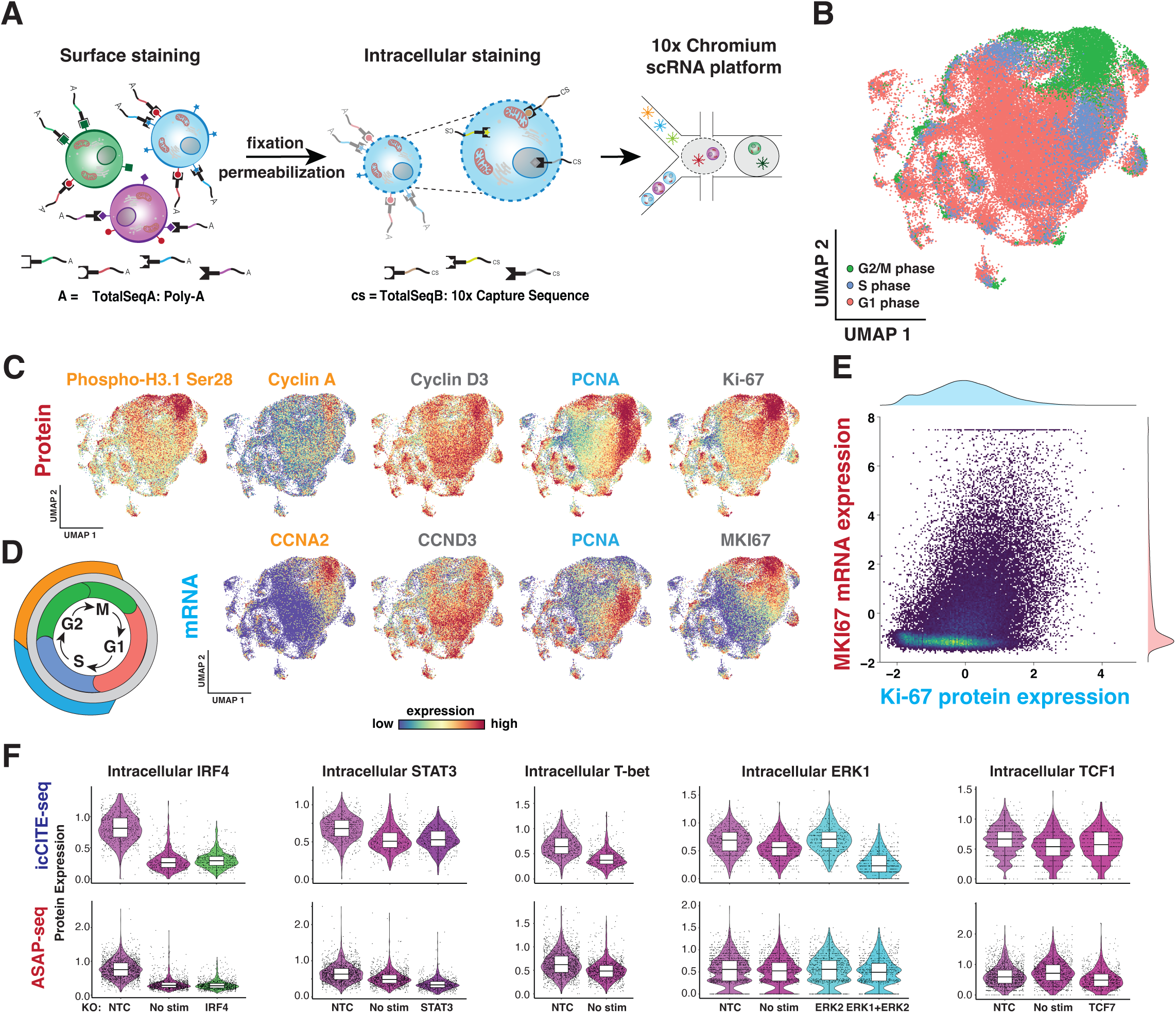
icCITE-seq enables simultaneous profiling of surface proteins, intracellular proteins and RNA levels at single-cell resolution. **a**, Schematic depicting the icCITE-seq workflow. Cells are first stained with TSA-conjugated antibodies targeting surface epitopes before fixation and permeabilization, followed by staining with TSB-conjugated antibodies targeting intracellular epitopes. **b**, icCITE-seq UMAP embedding of 57,776 single cells colored by cell cycle phase. **c**, UMAP embedding overlaid with protein expression (top) and RNA expression (bottom) of indicated protein and genes. **d**, Cell cycle schematic. Proteins in (**c**) are color-coded by their most prominent phase of expression in the cell cycle. **e**, Scatterplot with marginal histogram of MKI67 RNA pearson residuals vs CLR-normalized Ki-67 ADT counts. **f** Violin plots showing the distribution of CLR-normalized protein counts for indicated intracellular proteins and their associated perturbation condition in icCITE-seq (top) and ASAP-seq (bottom).

Compared to CITE-seq using fresh cells, fixed cells profiled by icCITE-seq exhibited on average, a 34% reduction in the number of unique molecular identifiers (UMIs) detected per cell and a 23% reduction in the number of detected genes (**Extended Data Fig. 1d**). However, this reduction was uniform and did not specifically affect genes from any particular condition (**Extended Data Fig. 1e**). Moreover, transcript-level detection from cells profiled using icCITE-seq and CITE-seq demonstrated a strong correlation in both non-stimulated and TCR-activated control cells, highlighting sensitivity of icCITE-seq across a broad range of detection of cellular states (ρ=0.95; ρ=0.93; **Extended Data Fig. 1f**). Notably, cells processed by icCITE-seq had significant loss in reads mapped to the mitochondrial genome, consistent with other previous reports using methanol-based fixatives (**Extended Data Fig. 1g**) ^9^.

Embedding of cells by icCITE-seq or CITE-seq RNA profiles alone largely separated cells by cell cycle state (G1, G2/M, S), confirming cell cycle stage as an important covariate to consider in subsequent analyses, which is consistent with previous reports (**Fig. 1b**) ^4,10,11^. To benchmark the specificity of intracellular staining, we next superimposed icCITE-seq intracellular protein abundances for PCNA, Ki-6, Cyclin D3, Cyclin A and phospho-histone H3.1 (Ser28) onto the icCITE-seq UMAP embedding (**Fig. 1c,d**). Reassuringly, this process yielded patterns which largely coincided with the expected cell cycle stage ^12–14^. Moreover, we found that the staining patterns were highly correlated with mRNA expression for the same genes as exemplified by Ki-67 (Spearman’s correlation = 0.425; *p*<2.2×10^-16^), further indicating that the staining was specific (**Fig. 1e**). Interestingly, we also noted a slight discrepancy between intracellular protein and mRNA expression levels for PCNA in cells under the G2/M phase, likely due to differential kinetics between RNA and protein expression.

To further validate the intracellular staining specificity in icCITE-seq, we compared intracellular protein tag signals with their respective CRISPR knockout in both icCITE-seq and ASAP-seq. While the distribution of IRF4, STAT3 and T-bet (*TBX21*) differed significantly between CRISPR-targeted and non-targeting controls or no stimulation controls in both ASAP-seq and icCITE-seq (**Fig. 1f**), robust staining of ERK2 and TCF7 could only be detected in icCITE-seq and ASAP-seq, respectively. We ascribe this disparity in antigen detection and specificity to the difference in fixation methods between the two assays. Importantly, we also observed several instances where staining signals for certain epitopes appeared biologically convincing but did not show a response in their corresponding knockout condition, suggesting non-specific staining (**Extended Data Fig. 1h**). These results highlight the critical need for thorough antibody validation and optimization to avoid confounding signals resulting from non-specific staining. Taken together, icCITE-seq enables nuanced annotation of cellular states by combining detection of surface and intracellular protein epitopes with RNA expression.

### Interrogation of T cell activation across multiple perturbations and modalities

To evaluate the biological effects resulting from our 107 targeted perturbations across chromatin accessibility, transcriptomic and protein modalities, we first minimized confounding effects from cell state on the transcriptome by regressing the signal using Seurat. We also removed contaminating CD8 T cells using unsupervised clustering and concomitant protein expression (**Supplementary Note**). We then embedded the resulting cells in joint Protein-ATAC and Protein-RNA space using two-modality weighted nearest neighbor (2WNN) UMAP ^15^ (**Fig. 2a,b**). Visualization of 2WNN embeddings of CD4 T cell single-cell chromatin accessibility and cell cycle-regressed RNA profiles revealed perturbation-specific chromatin and transcriptional changes, indicated by the formation of gRNA-specific clusters which flanked the central population (**Fig. 2a** and **Extended Data Fig. 2a-e**). For example, cells associated with gRNAs targeting in *IRF4*, *CHD4, BRD4*, or *CTCF* resolved into discrete clusters in UMAP embeddings for both chromatin accessibility and RNA, perhaps reflective of strong, divergent changes that manifested in both the epigenome and transcriptome. In contrast, cells with gRNAs targeting *SATB1*, *SETDB1* or *IKZF1* resulted in distinct clusters in ASAP-seq, but their effects were less prominent in RNA, suggesting that the gene expression changes associated with the perturbation of these genes were similar to that of other gRNAs.

**Figure 2.**
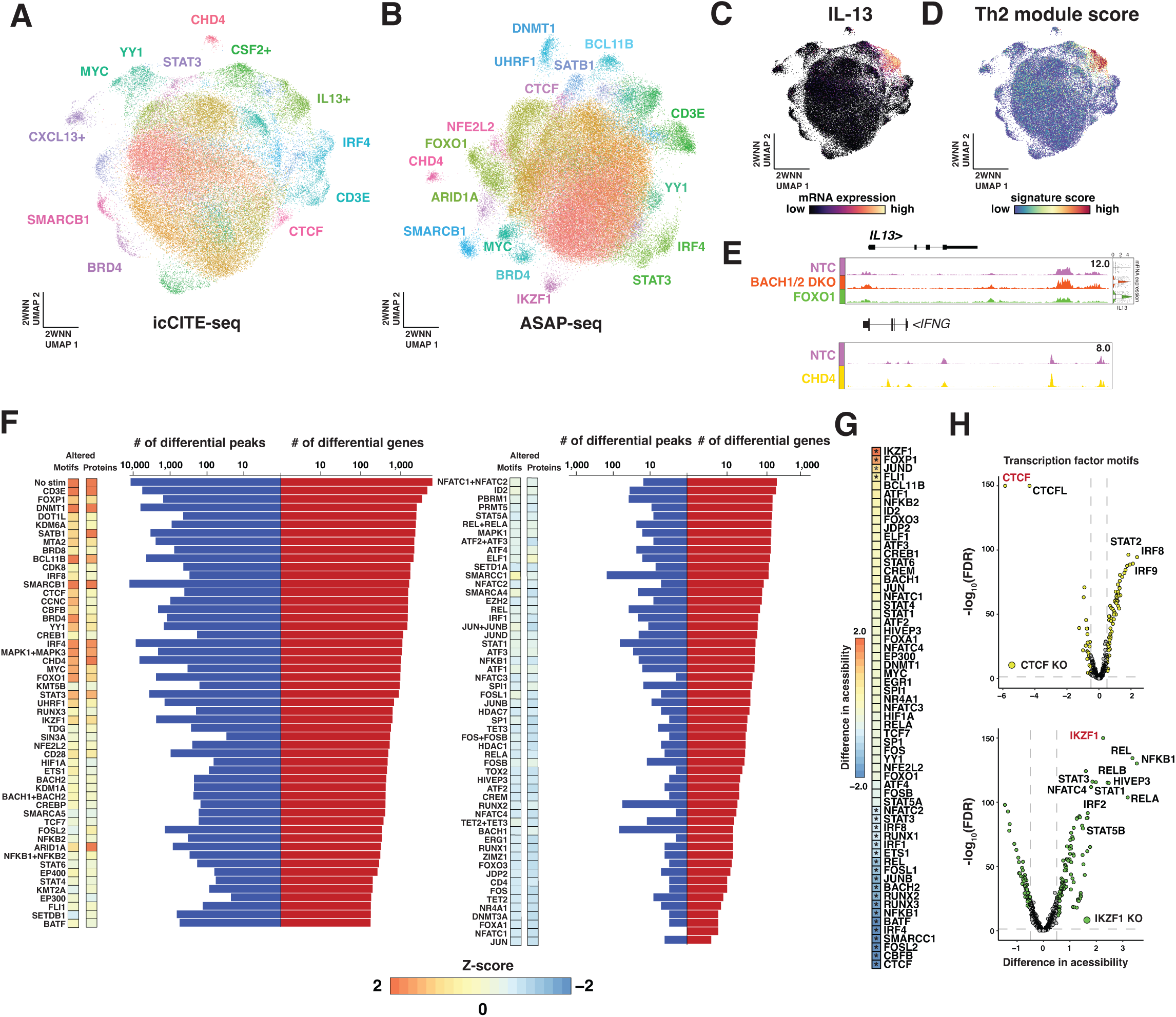
Multi-modal profiling of genetic perturbation phenotypes. **a**,**b** 2WNN UMAP embedding of the icCITE-seq (**a**) and ASAP-seq (**b**) dataset based on protein counts and cellular transcriptomes (**a**) or chromatin accessibility (**b**). Cells are colored by cluster assignment. **c**,**d** 2WNN UMAPs of joint single-cell mRNA expression and protein expression overlaid with IL13 mRNA expression (**c**) and Th2 module scores (**d**). **e**, Genomic tracks of *IL13* (top) and *IFNG* (bottom), indicating pseudo-bulk ATAC signal tracks across gRNAs with corresponding mRNA expression violin plots (top). **f**, The total number of significantly changed peaks and genes detected via ASAP-seq and icCITE-seq under each indicated knockout condition. Relative number of motif and protein changes are indicated. **g**, Heatmap representation of chromVAR bias-corrected transcription factor motif deviation scores for the indicated transcription factors and condition-matched knockout. **h**, Volcano plots showing transcription factor motifs with significantly changed chromatin accessibility profiles between NTC cells and knockouts targeting *CTCF* and *IKZF1* (FDR <= 0.05, chromVAR accessibility change >= 0.5).

We also noticed the presence of clusters that were not associated to a single gRNA in joint RNA-protein space, but were instead demarcated by the expression of genes such as *IL-13*, *CSF2* or *CXCL13* at the mRNA level (**Fig. 2a,c and Extended Data Fig. 3a,b**). Upon further examination, we found that this *IL-13*-expressing cluster (IC07) was phenotypically congruent with a classical helper Th2 signature and was enriched for gRNAs targeting *FOXO1*, *BACH2* and *BACH1/2* DKO, indicating that the specific perturbation of these genes may predispose towards a “Th2-like phenotype” upon TCR activation^16^ (**Fig. 2d and Extended Data Fig. 2c,d**). Visualization of aggregate pseudo-bulk chromatin accessibility at the *IL-13* locus revealed markedly changed chromatin profiles for *FOXO1*-depleted and BACH1/2-depleted cells when compared to non-targeting control cells (**Fig. 2e**). By contrast, *CHD4* KO cells exhibited elevated interferon signaling and T-bet expression accompanied with changes in accessibility at the *IFNG* locus (**Fig. 2e and Extended Data Fig. 3c,d**). Taken together, these observations highlight the utility of our multi-omic approach towards characterizing biological phenotypes resulting from targeted perturbations.

While previous studies coupling CRISPR perturbations with single cell readouts primarily focused on singular or dual modalities ^10,17–19^, our multi-omic approach enables the interrogation of CRISPR phenotypes across the epigenome, transcriptome and proteome. We therefore leveraged the multiple modalities of our dataset to systematically characterize the effects of genetic perturbations within our system (**Fig. 2f**). The number of chromatin accessibility regions altered by gene knockout was highly variable across perturbation conditions. For example, we observed 55 differentially accessible chromatin regions for *RUNX2* KO cells and 11,847 regions in *SMARCB1* KO cells (minimum log_2_ fold change gene expression 0.5 and *P*_adj._ <= 0.05). At the transcriptomic level, we found that *HIVEP3* KO cells significantly affected the expression of 21 genes, while *FOXP1* knockout affected the expression of 3,364 genes. Although the number of differentially accessible chromatin regions and number of differentially expressed genes were often correlated, we found notable exceptions in genes such as *MYC*, *SIN3A* and *ETS1*, where chromatin accessibility changes were relatively sparse despite the profound changes in the transcriptome. Collectively, these findings highlight the diverse mechanisms that govern gene regulation.

Eukaryotic gene regulation relies on the intimate cooperation between *trans*-acting factors and cis-regulatory DNA elements ^20,21^. However, this complex interplay comes in many forms with *trans*-factors often taking on activating, repressive or context-dependent roles, making mechanistic interpretation difficult. We therefore sought to extend our chromatin analyses by assessing accessible transcription factor motif deviations to quantify changes in transcription factor activities in response to *trans*-factor perturbation in the context of TCR signaling. As expected, we found that in general, the number of significantly altered TF motif accessibilities for a given perturbation was correlated with the number of differential chromatin accessibility regions (**Fig. 2f**). For the 61 out of the 107 perturbed *trans*-factors with known sequence motifs, we further queried whether or not the targeted perturbation significantly changed the accessibility at regions containing its motif, which could provide mechanistic insight on how the targeted *trans*-factor functions at their putative motifs. For example, consistent with its reported function, targeted perturbation of *CTCF* resulted in a significant decrease in chromatin accessibility at the corresponding CTCF motifs, indicating that CTCF facilitates the opening or maintenance of local chromatin at CTCF motif-occurring regions (median chromVAR accessibility loss, 5.86 and false discovery rate (FDR) < 0.05; **Fig. 2g,h**) ^22–24^. Similarly, abrogation of *IRF4* also resulted in a reduction of its motif accessibility (median chromVAR accessibility loss, 3.21 and FDR < 0.05; **Fig. 2g** and **Extended Data Fig. 3e**). In contrast, perturbation of *IKZF1* or *FOXP1* resulted in a significant increase in chromatin accessibility at their respective motifs (median chromVAR accessibility gain, 2.26 and FDR < 0.05, median chromVAR accessibility gain, 1.40 and FDR < 0.05, respectively; **Fig. 2g,h** and **Extended Data Fig. 3e**), suggesting that *IKZF1* and *FOXP1* may function as transcriptional repressors by decreasing chromatin accessibility at such regions. Notably, trans-factors of the AP-1 and Nuclear factor of activated T-cells (NFAT) transcription factor families including *FOS*, *JUN*, *NFATC3* and *NFATC4* did not display apparent local chromatin accessibility changes at their motifs upon perturbation, perhaps indicative of functional redundancy between other family members ^25–27^.

### Multi-omic mapping of perturbation-induced phenotypes at high-resolution

Recent CRISPR-based screening methods have been widely adopted to unbiasedly discover novel gene regulators ^28–30^, often by coupling with fluorescence activated cell sorting (FACS) to measure specific protein-based phenotypes. We reasoned that our dataset could be similarly used to simultaneously examine perturbation-induced phenotypes on specific mRNA transcripts, chromatin-based gene activity scores and protein markers by estimating the relative abundance of specific perturbation identities in single cells categorically binned as “high” or “low” expressers. As expected, applying this strategy to CD25 surface protein expression yielded well-known regulators of CD25 expression such as *CD3E*, *FOXP1* and *STAT5A* (**Fig. 3a,b**) ^31,32^. We also identified novel positive modulators of CD25 expression (e.g. *MYC*, *SMARCB1*, *CHD4*) as well as negative modulators such as *BACH1* and *BACH2*, which could also be similarly identified using *IL2RA* mRNA expression (**Extended Data Fig. 4a**). Furthermore, extending this analysis to ASAP-seq yielded highly consistent results, underscoring the robustness and applicability of our approach (ρ = 0.72; **Extended Data Fig. 4b,c**).

**Figure 3.**
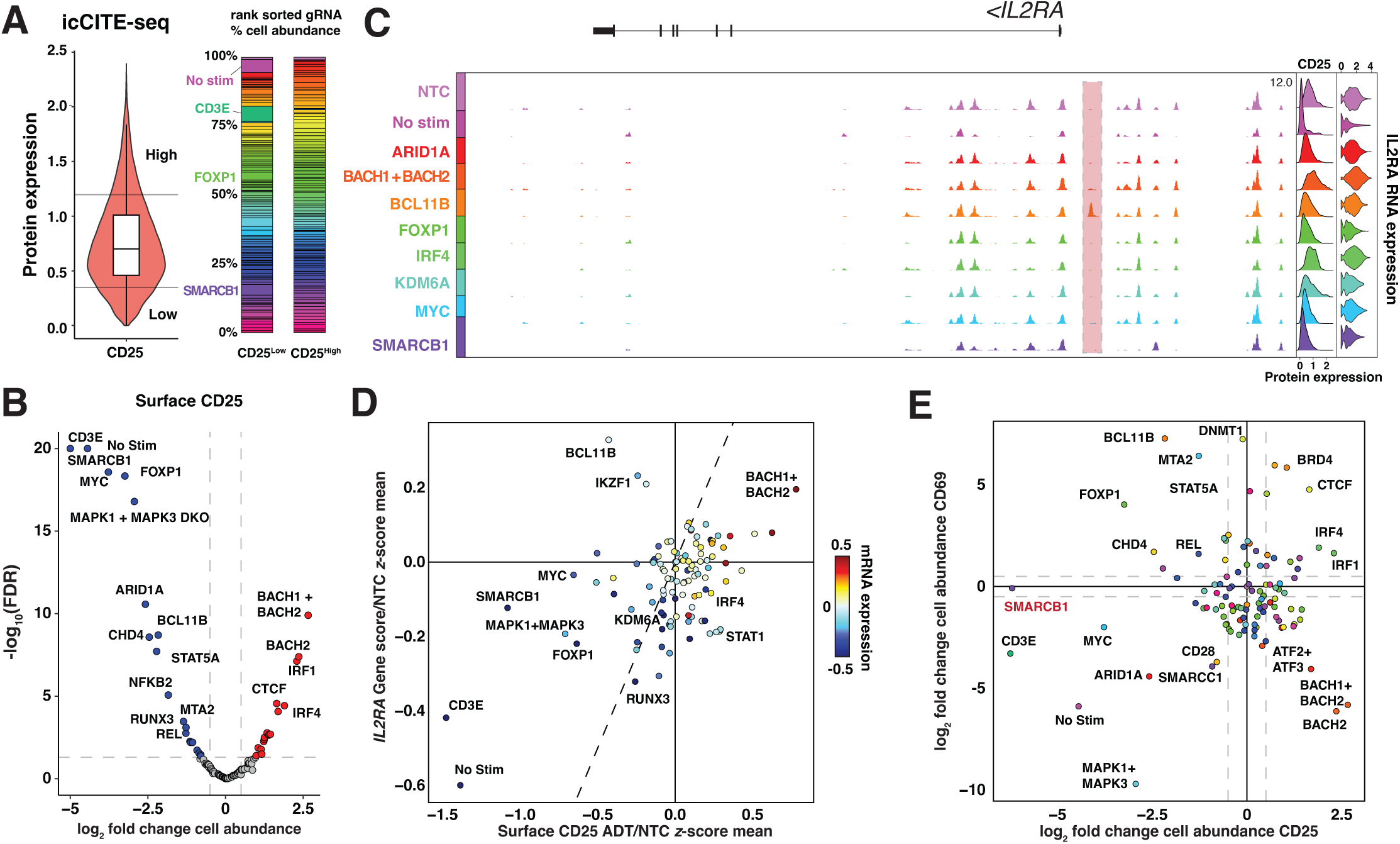
*in silico* screening identifies novel regulators of CD25 expression. **a**, Violin plot showing the distribution of CLR-normalized protein counts for CD25 (left), with upper and lower 15% cutoffs indicated. The boxplot spans from the first to the third quartile of the distribution, with the median positioned in the center. Whiskers represent the minimum and maximum values, excluding outliers. Stacked bar plots (right) depicting the proportional distribution of specific perturbation conditions in CD25^Low^ and CD25^High^ populations. Each color is representative of a specific knockout condition.**b**, Volcano plot depicting the log-fold change (LFC) in cell abundance across knockout conditions in CD25^Low^ versus CD25^High^ populations, relative to NTC cells. Significance threshold was set as FDR < 0.05 and absolute value of log_2_ odds ratio > 0.5. Statistical test was determined by a hypergeometric test and adjusted for multiple comparisons using the Benjamini-Hochberg approach. **c**, Genomic tracks of *IL2RA* (gene encoding CD25), indicating pseudo-bulk ATAC signal tracks across gRNAs with corresponding CLR-normalized protein abundance ridge plots and mRNA expression violin plots. A *BCL11B* knockout-specific differentially accessible region is highlighted in red. **d**, Scatterplot of mean gene activity scores for the *IL2RA* gene loci plotted against CLR-normalized mean CD25 protein tag counts associated with each knockout condition. Values are normalized against NTC cells. Color is indicative of mean *IL2RA* mRNA expression. **e**, Scatterplot showing the log-fold change (LFC) in cell abundance for CD25^Low^ versus CD25^High^ and CD69^Low^ versus CD69^High^ populations.

Examination of perturbation effects on chromatin accessibility at the *IL2RA* locus revealed a general concordance between the epigenome and mRNA/protein expression with a few striking exceptions such as *BCL11B*, which displayed a modest decrease in expression, despite a large gain of chromatin accessibility proximal to the *IL2RA* transcriptional start site (**Fig. 3c,d**). Interestingly, the specific perturbation of SWI/SNF-related chromatin remodeler *SMARCB1* resulted in abrogation of CD25 expression and was followed by extensive loss of TCR stimulation-induced open chromatin regions at the *IL2RA* locus, but unaffected CD69 expression, indicating decoupled regulation between these T cell effector genes (**Fig. 3c-e**). Lastly, we extended these analyses to the post-translationally modified intracellular epitope phospho-RPS6, which also identified *SMARCB1*, consistent with its role as a positive regulator of MTORC1 signaling (**Extended Data Fig. 4d**)^33^. Collectively, these results highlight the diverse regulatory circuitry that governs T cell activation.

### Single-cell analysis uncovers a heterogeneous response in *DNMT1*- and *UHRF1*-perturbed cells

In our icCITE-seq dataset, we observed the emergence of “Treg-like” cells in *DNMT1* and *UHRF1* perturbations, which is consistent with the notion that DNA methylation serves as one of the “gate-keepers” towards *FOXP3* expression (**Extended Data Fig. 5a-c**)^34–36^. However, this was a bifurcated response, which could be separated by Foxp3 protein and to a lesser extent, *FOXP3* mRNA expression, but not by *FOXP3* gene activity score (**Fig. 4a and Extended Data Fig. 5d**). Notably, Foxp3-negative cells were transcriptionally and epigenetically distinguishable from non-targeting control cells, which further indicated that this heterogeneous effect was not due to the presence of unperturbed cells (**Extended Data Fig. 5e**).

**Figure 4.**
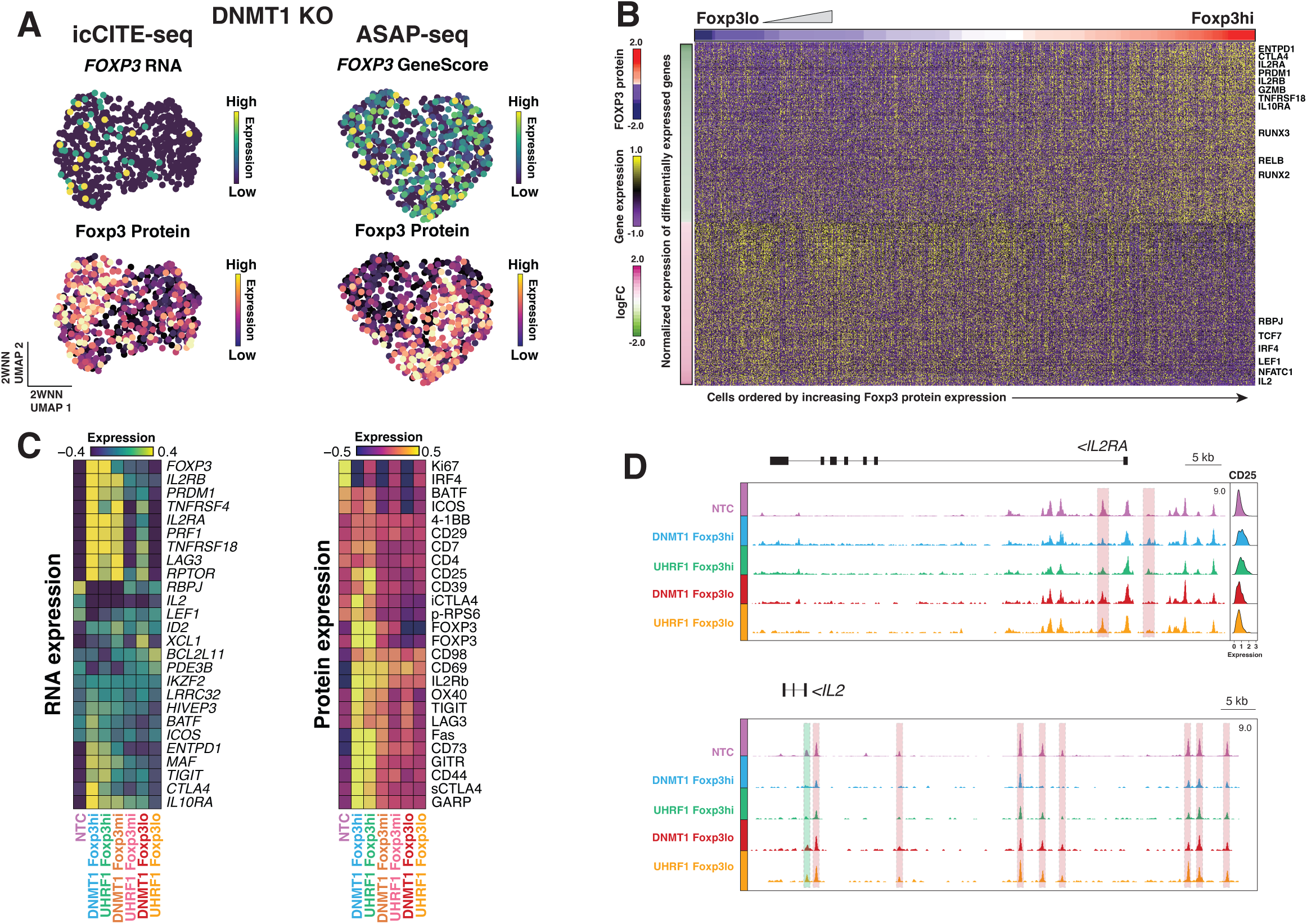
Single cell analysis resolves perturbation-induced heterogeneous cell states. **a**, 2WNN UMAP representations of *DNMT1* knockout cells in icCITE-seq (left) and ASAP-seq (right), overlaid with FOXP3 mRNA expression, gene accessibility and protein expression. **b**, Heatmap showing the differentially expressed genes in *DNMT1* and *UHRF1* knockout cells, comparing the top 15% versus the bottom 15% of cells based on FOXP3 protein expression levels. The *x* axis represents the ranked ordering of cells based on FOXP3 protein expression. **c**, Heatmap showing mean expression for mRNA expression (left) and protein expression (right) across indicated conditions. **d**, Genomic tracks of *IL2RA* (top) and *IL2* (bottom), indicating pseudo-bulk ATAC signal tracks across indicated conditions with corresponding CLR-normalized protein abundance ridge plots. Select differentially accessible regions are highlighted in red.

To examine how Foxp3 protein expression levels could putatively dictate specific transcriptional changes in these cells, we first quantified differential RNA and protein level changes across cells categorized with ‘high’ or ‘low’ Foxp3 expression levels (**Supplementary Note**). This analysis revealed an increase in expression of well-known Foxp3-inducible transcripts including *PRDM1*, *CTLA4*, *IL2RA*, *TNFRSF18*, *LAG3* and *IL10RA*, as well as a decrease in Foxp3-repressed genes such as *PD3EB* and *IL2* in cells expressing high levels of Foxp3 protein (**Fig. 4b,c**)*^37–39^*. Importantly, while the comparison of differential gene expression between bulk NTC and DNMT1/UHRF1 knockouts, as well as NTC and Foxp3^hi^ DNMT1/UHRF1 KO cells, showed strong correlation, the latter analysis demonstrated greater statistical power, identifying over 1,000 additional significantly changed genes, including critical genes such as *CTLA4*, *IL7R*, *CD44*, *STAT5A* and *IL10* (**Extended Data Fig. 5f,g**).

Examining these cells at the protein level, we noted concomitant increases in expression for GITR, CD25, LAG3 and CTLA4 (**Fig. 4c**). Interestingly, at the epigenetic level, we found that changes in chromatin accessibility were more varied and loci-specific (**Fig. 4d and Extended Data Fig. 5h**). For example, the *IL2RA* locus displayed relatively modest changes in chromatin accessibility despite clear increases in both RNA and protein (**Fig. 4c,d**). This was in contrast to *IL2*, where a sharp decrease in chromatin accessibility at the locus and nearby enhancer landscape was followed by a concomitant decrease in RNA expression. These observations support the notion that FOXP3 exploits various mechanisms, such as the co-binding of distinct and specific regulatory complexes, to regulate gene expression ^37^. Taken together, our results underscore the utility of examining the phenotypes of genetic perturbations at single-cell resolution across multiple modalities.

## Discussion

Here, we present icCITE-seq, a robust method that enables measuring surface and intracellular proteins alongside gene expression in thousands of single-cells. icCITE-seq enables profiling of cytoplasmic, nuclear, and phosphorylated intracellular epitopes with minimal loss of RNA integrity, enhancing our ability to examine diverse cellular processes while providing a more holistic view of intracellular signaling and cellular dynamics. To validate the specificity of intracellular antibody signals in icCITE-seq, we profiled the assay with arrayed CRISPR knockouts and demonstrated that most detected signals accurately corresponded to their targeted intracellular epitopes, confirming the reliability of the approach.

By integrating genetic perturbation data from icCITE-seq and ASAP-seq, we fine-mapped the phenotypic responses of over 100 nuclear factors in the context of TCR stimulation across three distinct modalities. Our results revealed the diverse mechanisms that coordinate TCR signaling responses across chromatin, transcriptional, and protein levels. By quantifying intracellular proteins, we were able to disentangle heterogeneous cellular states, as exemplified by FOXP3 in *DNMT1* and *UHRF1* knockouts (**Fig. 4**). Additionally, leveraging these intracellular protein signals enhanced the sensitivity of differential gene expression analysis. Our framework therefore demonstrates how integrating genetic perturbations with intracellular protein profiling can provide valuable insights into cellular complexity. We anticipate that the expansion of available intracellular antibody panels will further enhance our understanding of signaling pathways across diverse biological contexts.

For future applications, we emphasize that care should be taken with respect to biological conclusions, which may require orthogonal validation methods. Indeed, we noticed that some signals such as phospho-NRF2 and HIF1α were non-specific (**Extended Data Fig. 1h**), emphasizing the need for rigorous antibody selection and validation to ensure accurate profiling. In total, our results indicate that icCITE-seq is a powerful tool for understanding the behavior of single cells in complex settings.

## Methods

### Flow Cytometry

For flow cytometric analysis, cells were stained with appropriate antibodies for cell surface proteins and Live/Dead dye. Cells were fixed and permeabilized using the Foxp3/Transcription Factor Staining Buffer Set (Thermo Fisher), followed by intracellular staining for FOXP3 (Invitrogen; clone 236A/E7). Stained cells were analyzed or sorted on the BD FACSAria fusion system.

### Arrayed Cas9 Ribonucleotide Protein Preparation and Electroporation

Cas9 ribonucleotide protein (RNP) preparation was performed essentially as previously described ^5^. Briefly, lyophilized cRNAs and tracrRNAs (purchased from IDT) were reconstituted to 400 µM and mixed at a 1:1 v/v ratio. The resulting mixture was heated at 95°C for 5 minutes, followed by incubation for 15 minutes at room temperature to complex the gRNAs. For the 70-plex and 100-plex knockout experiments, two gRNAs targeting the same gene were pooled together to maximize knockout efficiency ^40^. Subsequently, 30 µg Cas9 protein (TakaraBio, Cat#Z2640N) was added to each well and mixed by pipetting, followed by incubation at room temperature for an additional 15 minutes. 12.7 µL of the resulting RNP complexes were mixed with 1E6 cells suspended in 20 µL Lonza P2 primary nucleofection buffer and then transferred into a 16-well electroporation cuvette plate (Lonza, Cat#V4XP-2032). Cells were pulsed with the EH100 program and immediately supplemented with 100 µL pre-warmed T cell culture medium. Electroporated cells were then incubated at 37°C for 10 minutes before transfer into 96-well U-bottom plates for culture, supplemented with 500 IU/mL IL-2. A list of all gRNAs used in this study can be found in **Supplementary Table 1**.

### Human CD4 T Cell Multiplexed Perturbations

Human naive CD4 T cells were enriched from PBMC by magnetic negative selection using the EasySep™ Human Naïve CD4+ T Cell Isolation Kit II (STEMCELL Technologies, Cat#17555) as per manufacturer’s instructions. After enrichment, cells were cultured in T cell culture medium consisting of RPMI supplemented with 10% Fetal Bovine Serum, 10 mM HEPES, 2 mM GlutaMax (Gibco, Cat#35050-061), 1x MEM Non-Essential Amino Acids (Gibco, Cat#11140-050), 1 mM Sodium pyruvate, 55 µM 2-mercaptoethanol and 100 IU/mL IL-2 at a density of 1E6 cells/mL and stimulated with anti-human CD3/CD28 Dynabeads (ThermoFisher, Cat#11131D) with a 1:1 cells-to-beads ratio. After stimulation (72 hours), beads were removed and the cells were further expanded for five days in T cell culture medium supplemented with IL-2, while maintaining a density of 1E6 cells/mL. Expanded cells were electroporated with Cas9 ribonucleoprotein (RNP) complex and then further expanded in T cell culture medium with 500 IU/mL IL-2. On day 15, cells were re-stimulated with anti-human CD3/CD28 Dynabeads (ThermoFisher, Cat#11132D), supplemented with 100 IU/mL IL-2 for 72 hours.

### Cell surface staining with barcoded antibodies

72 hours post-stimulation, magnetic beads were removed and the perturbed cells were stained with defined combinations of barcoded TotalSeq-A and TotalSeq-B conjugated hashing antibodies (BioLegend; see **Supplementary Table 2** for a list of antibodies). Briefly, up to 1 million cells per sample were resuspended in Cell Staining Buffer (BioLegend, Cat#420201) and stained with 1/100 dilutions of each (combinations of two) TotalSeq-A and TotalSeq-B hashtag antibody and 1/1000 dilution of LIVE/DEAD™ Fixable Near-IR Dead Cell Stain Kit (Invitrogen, Cat#L10119) for 30 minutes on ice. Subsequently, cells were washed 3x with 1 mL of Cell Staining Buffer. After the final wash, cells were resuspended with 200 µL Cell Staining Buffer and filtered through a 70 µM cell strainer before live cell sorting on a BD FACSAria-III (Becton Dickinson). Approximately 1.5 million sorted cells were pooled and resuspended in Cell Staining Buffer and incubated for 10 minutes with TruStain FcX (BioLegend, Cat#4223020). Cells were then surface stained with the indicated TotalSeq-A antibody cocktails for 30 minutes at 4°C, as recommended by the manufacturer (BioLegend) and then washed 3x with 1 mL of Cell Staining Buffer. After the final wash, cells were resuspended in PBS for downstream processing with CITE-seq using the 10x Chromium controller with the Chromium Next GEM Single Cell 3’ Kit v3.1 (10X Genomics, Cat#1000268) and 3’ Feature Barcode Kit (10X Genomics, Cat#1000262) or ASAP-seq with the Chromium Next GEM Single Cell ATAC Library & Gel Bead Kit v1.1 (10X Genomics, Cat#1000175) as previously described^1,5^.

### Intracellular ASAP-seq

After surface staining with TotalSeq-A barcoded antibodies as described in the cell surface staining methods section, approximately 1 million cells were resuspended with Cell Staining Buffer and then pelleted by centrifugation for five minutes at 500xg. The supernatant was carefully removed and the resulting cell pellet was resuspended using the residual liquid volume. Subsequently intracellular staining for ASAP-seq was performed with fixed and permeabilized cells as previously described^5^.

### Intracellular CITE-seq (icCITE-seq)

After surface staining with TotalSeq-A barcoded antibodies as described in the cell surface staining methods section, approximately 1 million cells were resuspended with Cell Staining Buffer and then pelleted by centrifugation for five minutes at 500xg. The supernatant was carefully removed and the resulting cell pellet was resuspended using the residual liquid volume. While gently vortexing the cells with a benchtop vortex, the cells were fixed and permeabilized by the dropwise addition of 1 mL pre-chilled True-Phos™ Perm Buffer (BioLegend, Cat#425401) and then incubated at -20°C overnight.

The next day, cells were pelleted by centrifugation at 4°C at 2000xg for five minutes and washed with ice-cold 2 mL of Intracellular Wash Buffer (1x) with 2 mM DTT. The centrifugation step was repeated once to completely remove the supernatant and then intracellular staining was performed using Intracellular Wash Buffer (1x) (BioLegend, custom part no. 900002577), with the addition of 2 mM DTT, TruStain FcX (BioLegend, Cat#4223020), True Stain Monocyte Blocker (BioLegend, Cat#426102) and 1U/uL Protector RNase Inhibitor (Roche, Cat#3335402001), as per manufacturer’s recommendations. After staining, cells were washed 3x with 1 mL of Intracellular Wash Buffer (1x) and then resuspended in Intracellular Wash Buffer (1x) with 2 mM DTT and 0.2U/uL RNase inhibitor. A maximum of 4 uL of the resulting cell suspension was used for processing with the Chromium Next GEM Single Cell 3’ Kit v3.1 (10X Genomics, Cat#1000268) and 3’ Feature Barcode Kit (10X Genomics, Cat#1000262) according to the manufacturer’s protocols.

### Data availability

Raw sequencing data for icCITE-seq and ASAP-seq are deposited on the DNA Data Bank of Japan Sequence Read Archive, DDBJ: DRAXXXXXX.

## Acknowledgements

We thank the members of the Sakaguchi laboratory for technical advice and discussions. This research was supported by Grants-in-Aid by the Japan Society for Promotion of Science for Specially Promoted Research no. 16H06295 and the Japan Agency for Medical Research and Development for Leading Advanced Projects for Medical Innovation.

## Author contributions

K.Y.C., Y.T. and S.S. conceived and designed experiments with input from A.Z.F., J.C. and K.T. K.Y.C., Y.T., K.M., T.K., M.H., and K.I. carried out experiments. K.Y.C., Y.T. and K.M. carried out analyses. K.Y.C. conceived and designed the methods with input from A.Z.F., J.C. and K.T. S.S. supervised the work. K.Y.C., Y.T. and S.S. drafted the manuscript with input from all other authors.

## Author information

Correspondence and requests for materials should be addressed S.S.

## Supplementary Information

Supplementary Note: Supplementary Methods

## Supplemental Tables

Supplementary Table 1: gRNAs used in this study.

Supplementary Table 2: Antibodies used in this study.

**Extended Data Figure 1.**
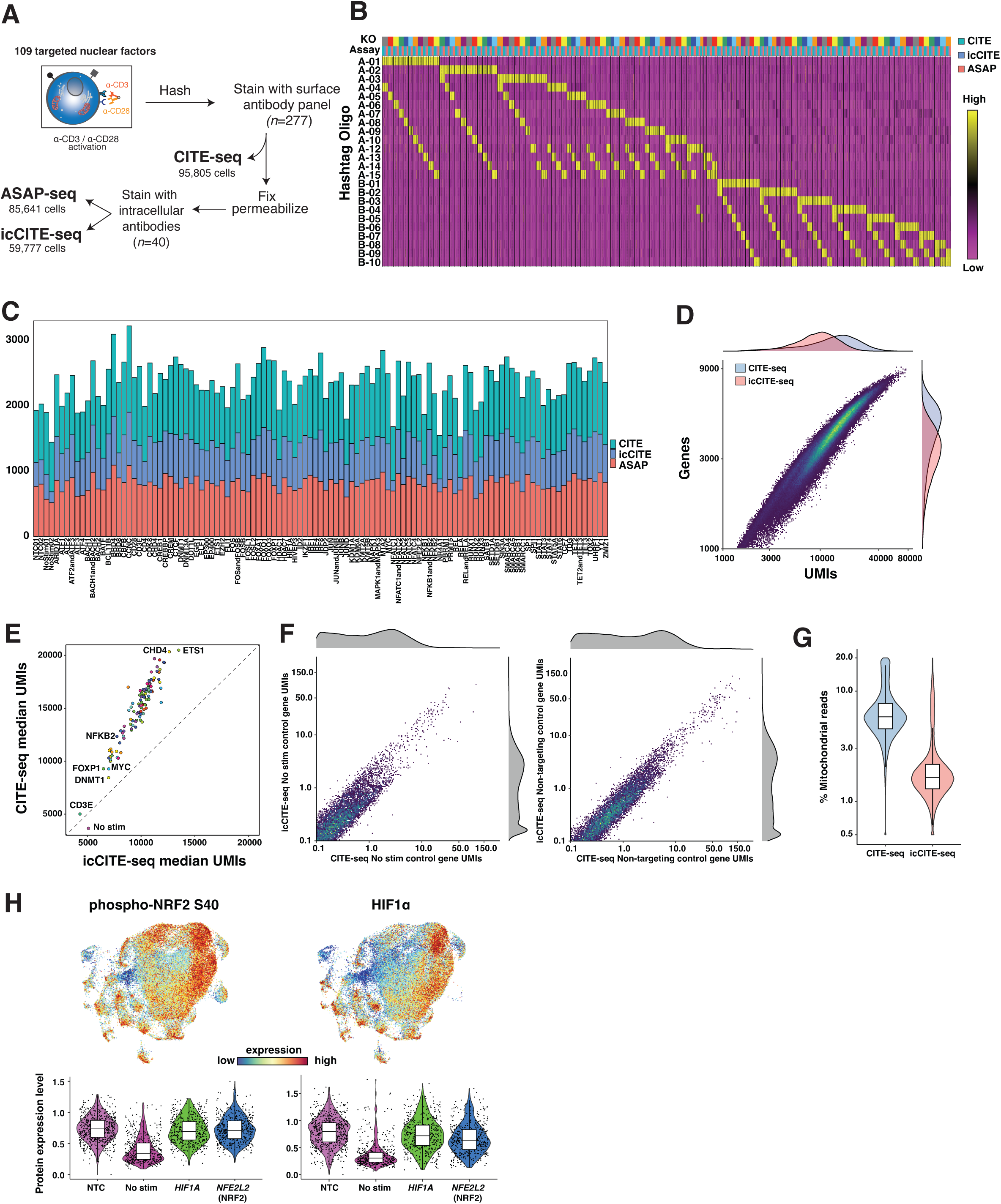
Additional benchmarking of the icCITE-seq workflow. **a**, Schematic of experimental design. CRISPR-edited cells are first stained with oligo-conjugated hashtag antibodies and then pooled for processing by CITE-seq, ASAP-seq and icCITE-seq. gRNA identities are demultiplexed using hashing antibody counts. **b**, Heatmap of cell demultiplexing with hashing antibodies, indicating normalized abundance of each hashtag. TotalSeqA^TM^ and TotalSeqB^TM^ hashtag antibodies are denoted as A- or B-, respectively. **c**, Stacked barplot showing the distribution of cell counts of demultiplexed knockout conditions across CITE-seq, icCITE-seq and ASAP-seq. **d**, icCITE-seq RNA library complexity compares well to scRNA-seq. Scatterplot of number of transcripts (UMIs) (*x* axis) and genes (*y* axis) detected in CITE-seq and icCITE-seq. Histograms on the top (for UMIs) and right (for genes) show marginal distributions. **e**, Correlation analysis of median transcripts detected in icCITE-seq (*x* axis) and CITE-seq (*y* axis), stratified by knockout condition. **f**, Correlation plot depicting gene-level transcript (UMI) detection in CITE-seq (*x* axis) and icCITE-seq (*y* axis) in non-stimulated control cells (left) and non-targeting control cells (right). **g**, Violin plot depicting percentage of mitochondrial genes detected in CITE-seq and icCITE-seq. The boxplot spans from the first to the third quartile of the distribution, with the median positioned in the center. Whiskers represent the minimum and maximum values, excluding outliers. **h**, UMAP embedding overlaid with expression of two intracellular protein markers (top). Violin plots showing the distribution of CLR-normalized protein counts for indicated intracellular proteins and perturbation conditions in icCITE-seq (bottom).

**Extended Data Figure 2.**
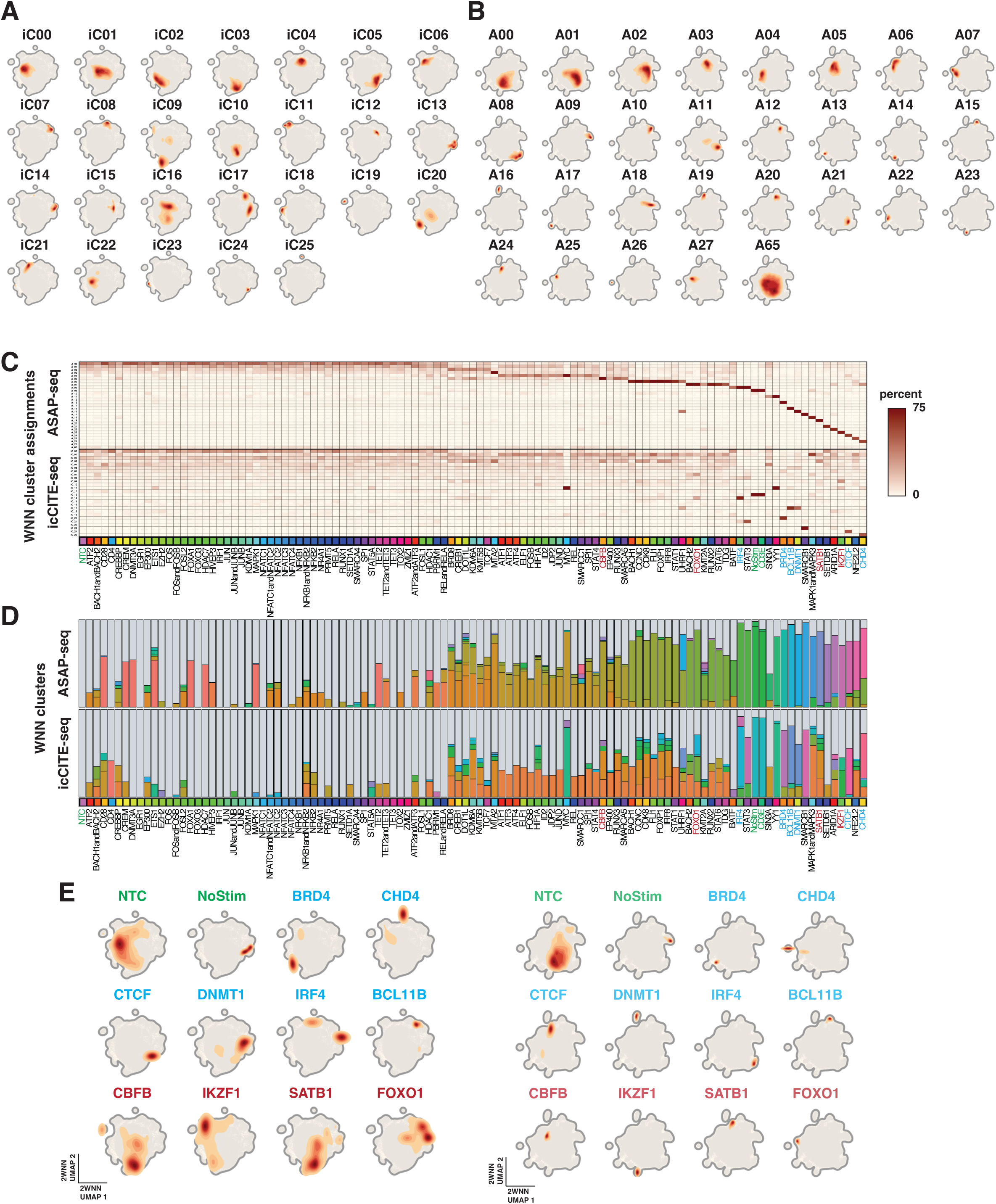
Additional benchmarking of the icCITE-seq workflow. **a**,**b** Visualization of cluster assignments in icCITE-seq (**a**) and ASAP-seq (**b**) by density plots. **c**, Heatmaps illustrating the percentage distribution of cells across each 2WNN cluster for ASAP-seq (top) and icCITE-seq (bottom) under 109 distinct knockout conditions. **d**, Stacked barplots representing the percent distribution of cells across 2WNN clusters under 109 distinct knockout conditions for ASAP-seq (top) and icCITE-seq (bottom). Each color corresponds to a specific 2WNN cluster. **e**, Density plots showing the spatial distribution of cells in the 2WNN UMAP embedding for icCITE-seq (left) and ASAP-seq (right) datasets under indicated knockout conditions.

**Extended Data Figure 3.**
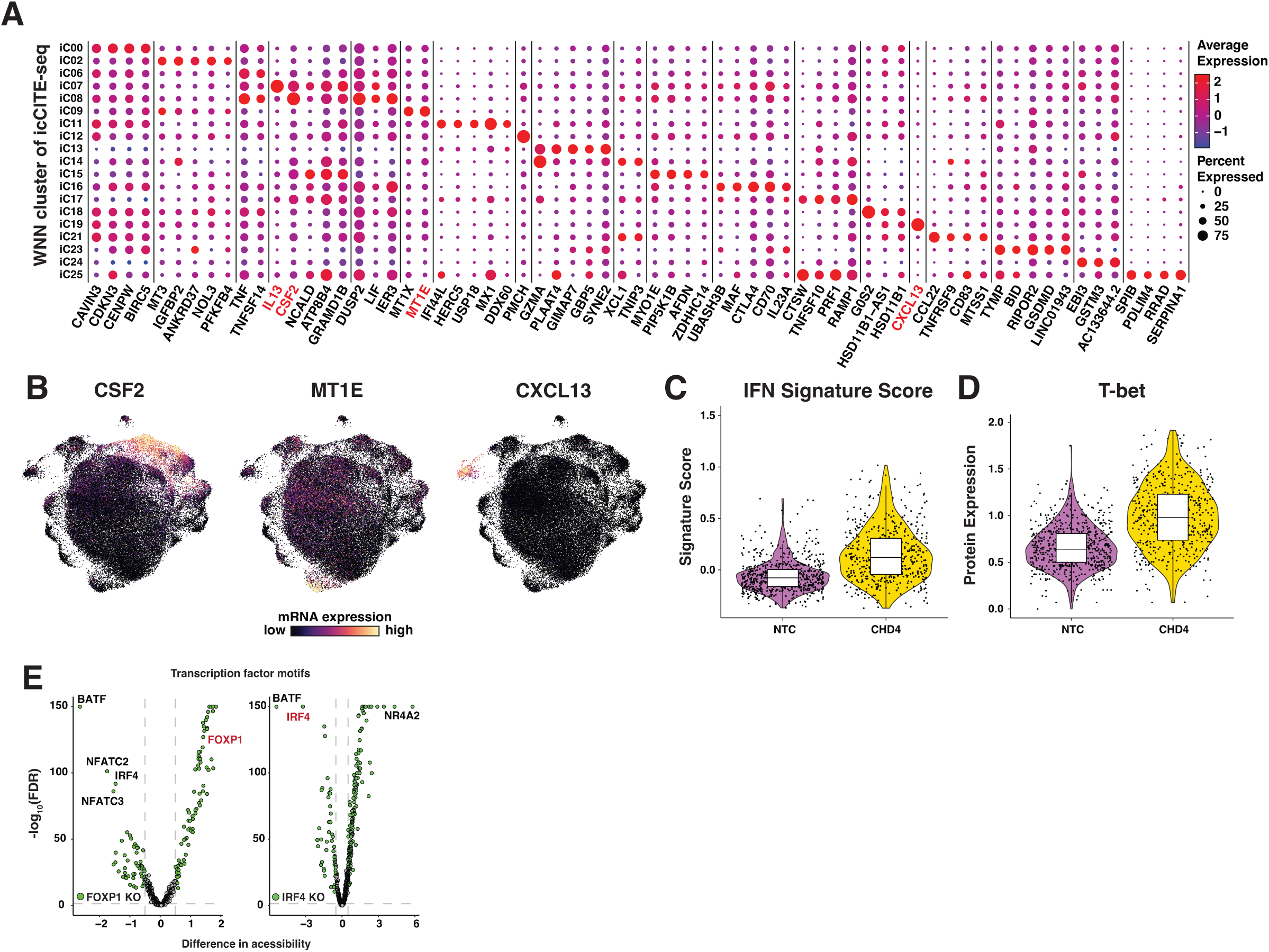
Additional phenotyping for perturbation conditions. **a**, Expression levels of selected RNA markers across identified WNN clusters. The dot size indicates the percentage of cells expressing the gene within each cluster and color is representative of the average expression level. **b**, 2WNN UMAPs of joint single-cell mRNA expression and protein expression overlaid with the expression of the indicated genes. **c**,**d** Violin plots depicting the IFN signature score (**c**) and CLR-normalized intracellular T-bet protein expression (**d**) between NTC and CHD4 knockout cells. **e**, Volcano plots showing transcription factor motifs with significantly changed chromatin accessibility profiles between NTC cells and knockouts targeting *FOXP1* and *IRF4* (chromVAR accessibility change >= 0.5 and FDR <= 0.05).

**Extended Data Figure 4.**
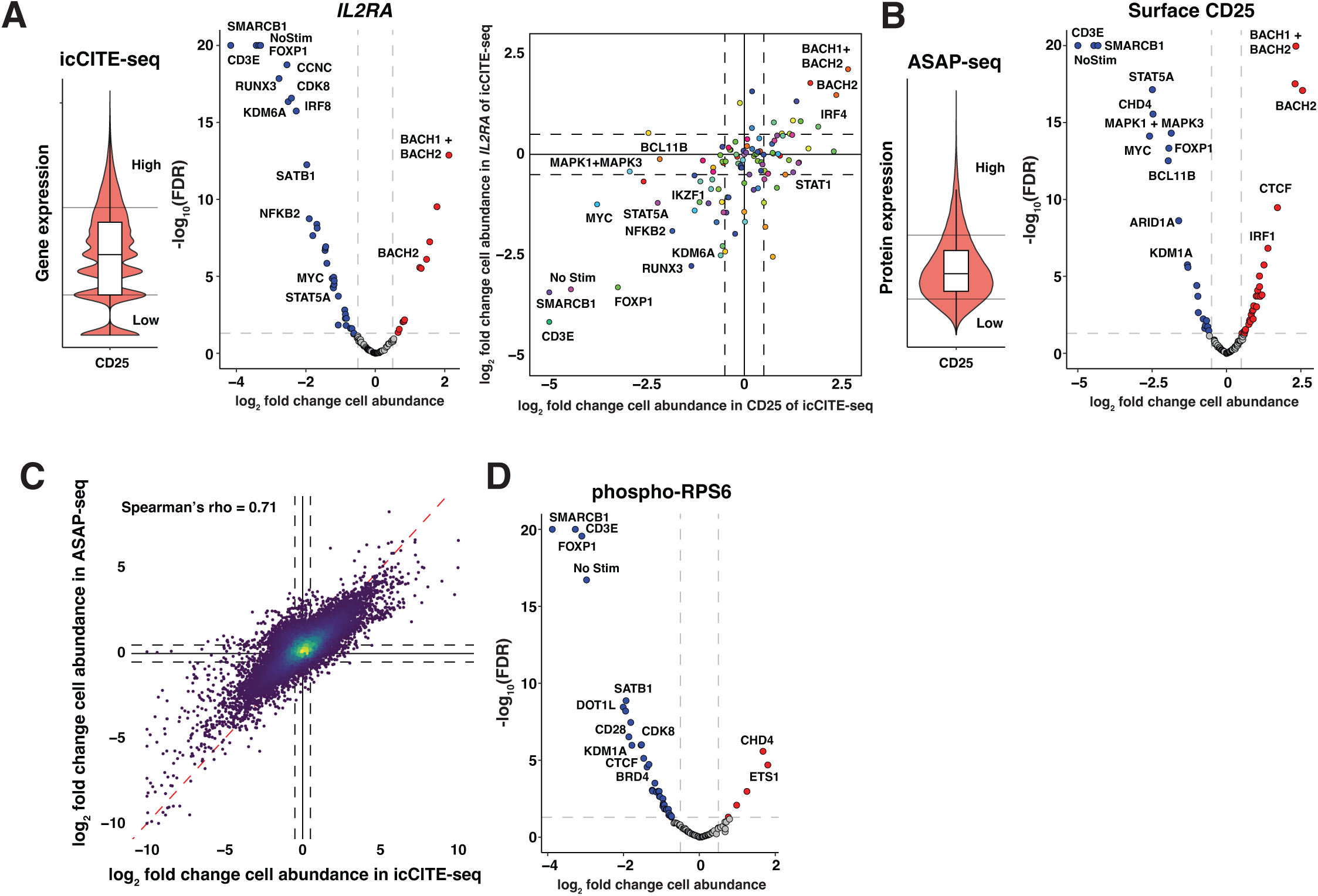
Additional analyses for in silico CRISPR screening. **a**, Violin plot showing the distribution of gene expression for *IL2RA* (left). Upper and lower 15% cutoffs are indicated. Volcano plot depicting the log-fold change (LFC) in cell abundance across knockout conditions in *IL2RA*^Low^ versus *IL2RA*^High^ populations, relative to NTC cells (middle). Correlation analysis of log-fold change cell abundance in CD25 protein and IL2RA gene expression (right). **b**, Analyses related to (**a** and Fig. 3a), but using CD25 protein as assessed by ASAP-seq. **c**, Correlation analysis of log-fold change cell abundance across 139 different proteins and 108 knockout conditions as assessed by icCITE-seq (*x* axis) and ASAP-seq (*y* axis). **d**, Volcano plot depicting the log-fold change (LFC) in cell abundance across knockout conditions in phospho-RPS6^Low^ versus phospho-RPS6^High^ populations, relative to NTC cells. Boxplots span from the first to the third quartile of the distribution, with the median positioned in the center. Whiskers represent the minimum and maximum values, excluding outliers.

**Extended Data Figure 5.**
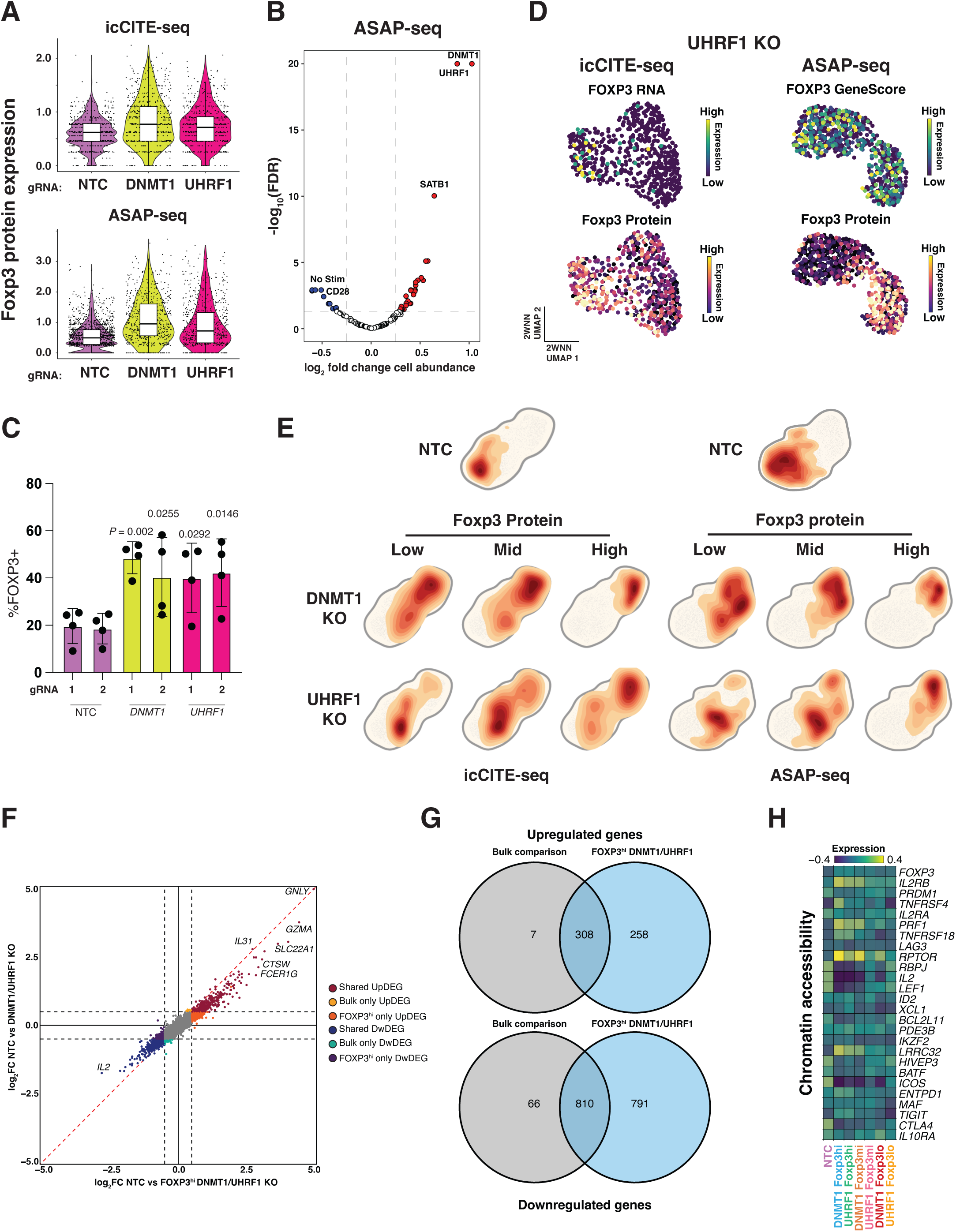
Supporting analyses for *DNMT1*/*UHRF1* knockouts. **a**, Violin plots showing the distribution of CLR-normalized protein counts for FOXP3 intracellular proteins under the indicated conditions in icCITE-seq (top) and ASAP-seq (bottom). **b**, Volcano plot depicting the log-fold change (LFC) in cell abundance across knockout conditions in FOXP3^Low^ versus FOXP3^High^ populations, relative to NTC cells. **c**, Analysis of FOXP3+ Treg cells after TCR stimulation in CD4+ T cells with indicated genetic perturbations. **d**, 2WNN UMAP representations of *UHRF1* knockout cells in icCITE-seq (left) and ASAP-seq (right), overlaid with FOXP3 mRNA expression, gene accessibility and protein expression. **e**, Density plots showing the spatial distribution of cells in the 2WNN UMAP embedding for icCITE-seq (left) and ASAP-seq (right) datasets under indicated knockout conditions. **f**, Scatter plot of gene expression fold changes from NTC versus FOXP3^hi^ DNMT1/UHFR1 KO (*x* axis) and bulk NTC versus DNMT1/UHRF1 KO (*y* axis). **g**, Venn diagram depicting overlap of differentially upregulated (top) and downregulated (bottom) genes in Bulk versus FOXP3^hi^ comparisons, related to (**f**). **h**, Heatmap showing mean gene scores for different loci across indicated conditions.

